# Probing Specificities of Alcohol Acyltransferases for Designer Ester Biosynthesis with a High-Throughput Microbial Screening Platform

**DOI:** 10.1101/2021.06.26.450049

**Authors:** Jong-Won Lee, Hyeongmin Seo, Caleb Young, Cong T. Trinh

## Abstract

Alcohol acyltransferases (AATs) enables microbial biosynthesis of a large space of esters by condensing an alcohol and an acyl CoA. However, substrate promiscuity of AATs prevents microbial biosynthesis of designer esters with high selectivity. Here, we developed a high-throughput microbial screening platform that facilitates rapid identification of AATs for designer ester biosynthesis. First, we established a microplate-based culturing technique with *in situ* fermentation and extraction of esters. We validated its capability in rapid profiling of the alcohol substrate specificity of 20 chloramphenicol acetyltransferase variants derived from *Staphylococcus aureus* (CAT_Sa_) for microbial biosynthesis of acetate esters with various exogeneous alcohol supply. By coupling the microplate-based culturing technique with a previously established colorimetric assay, we developed a high-throughput microbial screening platform for AATs. We demonstrated that this platform could not only confirm CAT_Sa_ F97W with enhanced isobutyl acetate synthesis but also identify three ATF1_Sc_ (P348M, P348A, and P348S) variants, derived from *Saccharomyces cerevisiae*’s AAT and engineered by model-guided protein design, for enhanced butyl acetate production. We anticipate the high-throughput microbial screening platform is a useful tool to identify novel AATs that have important roles in nature and industrial biocatalysis for designer bioester production.

## 1. INTRODUCTION

Esters are an important class of industrial chemicals with broad applications as flavors, fragrances, cosmetics, pharmaceuticals, green solvents, and advance biofuels (Lee and Trinh 2020). Currently, esters are mainly produced by chemical synthesis from petroleum-based feedstocks that are neither renewable nor sustainable (Seo et al. 2019). Alternatively, microbial conversion has emerged as an alternative route for renewable and sustainable production of esters (Layton and Trinh 2014; Layton and Trinh 2016a; Layton and Trinh 2016b; Lee and Trinh 2019; Rodriguez et al. 2014; Tai et al. 2015b). Critical to the microbial biosynthesis of esters is the requirement of alcohol acyltransferases (AATs, EC 2.3.1.84) that catalyze a thermodynamically favorable condensation of an alcohol and acyl-CoA in an aqueous environment. Since esters are commonly found in nature such as fruits (e.g., apple (Li et al. 2006; Souleyre et al. 2014; Souleyre et al. 2005), apricot (Gonzalez-Aguero et al. 2009), banana (Beekwilder et al. 2004), melon (El-Sharkawy et al. 2005; Lucchetta et al. 2007), papaya (Balbontin et al. 2010), and strawberry (Aharoni et al. 2000; Beekwilder et al. 2004; Cumplido-Laso et al. 2012; Gonzalez et al. 2009)) or yeast fermentation (Tai et al. 2015b; Verstrepen et al. 2003), various eukaryotic AATs have been identified and recently exploited for microbial biosynthesis of esters using synthetic biology and metabolic engineering approaches (Layton 2014; Lee and Trinh 2018; Rodriguez et al. 2014; Tai et al. 2015a). Recent discovery and repurposing of prokaryotic chloramphenicol acetyltransferases (CATs, EC 2.3.1.28) to function as AATs have further expanded the library of ester-producing enzymes (Rodriguez et al. 2014; Seo et al. 2019). However, due to substrate promiscuity of these AAT/CAT enzymes, controllable microbial synthesis of designer esters with high selectivity remains a significant challenge (Layton and Trinh 2014; Lee and Trinh 2019).

Bioprospecting and protein engineering are promising strategies to find novel AATs with high specificity and activity toward a target ester. For instance, AAT of *Actinidia chinensis* (AAT_Ac_) was engineered to create a AAT_Ac_ S99G variant that enhanced butyl octanoate production in *Escherichia coli* about 4.5-fold higher than the wildtype (Chacon et al. 2019). Similarly, Seo *at al*. reported that a single F97W mutation in CAT of the mesophilic *Staphylococcus aureus* (CAT_Sa_), identified by a model-guided protein design, achieved ∼3.5-fold increase in isobutyl acetate (IBA) production in a thermophilic, cellulolytic bacterium *Clostridium thermocellum* (Seo et al. 2019). By combining both bioprospecting and model-guided protein engineering strategies, novel CATs have recently been discovered with improved efficiency, robustness, and compatibility (Seo et al. 2020). Even though research efforts in identifying beneficial AATs/CATs with high specificity and activities are promising, large space of novel AATs/CATs are still underexplored.

To access the specificities and activities of AATs/CATs directly, the enzymes need to be purified and characterized. Two colorimetric assays have been developed to determine AAT/CAT activities including the 5,5′-dithiobis-(2-nitrobenzoic acid) (DTNB) assay (Kruis et al. 2017; Seo et al. 2019; Tai et al. 2015b) and the α-ketoglutarate dehydrogenase (α-KGDH)-coupled assay (Knight et al. 2014; Lin et al. 2016). These assays are designed to quantify free CoAs released from the AAT/CAT esterification of alcohols and acyl-CoAs by measuring either the 412 nm absorbance of yellowish 5-thio-2-nitrobenzoic acid (TNB) for the DTNB assay or the 340 nm absorbance of nicotinamide adenine dinucleotide (NADH) for (α-KGDH)-coupled assay. The key advantage of direct AAT/CAT measurement is that the assays can be performed in a high-throughput manner; however, some disadvantages for screening a large space of AATs/CATs include requirement of expensive acyl-CoA reagents and enzyme purification. Alternatively, direct measurement of esters for rapid, high-throughput screening of AAT/CAT specificities and activities *in vivo* can be attractive before determining the catalytic efficiencies in depth for promising enzyme candidates. Here, esters produced by microorganisms can be extracted with a solvent (e.g., hexane or hexadecane) and measured in a separate step. While the conventional gas chromatography coupled with mass spectrometer (GC/MS) is accurate in identifying and quantifying esters, it is low-throughput and expensive. Fortunately, the colorimetric assay, based on the hydroxylamine/iron chemistry, can rapidly quantify esters in a high-throughput manner by first generating the ferric hydroxamate via the two steps of chemical reactions and then measuring its absorbance at 520 nm (Hill 1946; Lobs et al. 2016; Stern and Shapiro 1953; Wofford et al. 1986).

In this study, we aimed to develop a high-throughput microbial screening platform to identify novel AATs/CATs for designer ester biosynthesis in a simple, rapid, and efficient manner. We started by establishing a microplate-based culturing technique with *in situ* fermentation and extraction of esters. By coupling the microplate-based culturing technique with a modified colorimetric assay, we developed a high-throughput microbial screening platform to identify novel AATs. The platform can measure both esters and cell growth, which helps not only screen relative AATs/CATs specificities and activities rapidly but also evaluate the effect of expressing these enzymes on microbial health. We validated the developed high-throughput microbial screening platform by probing the alcohol substrate preference of 20 engineered CAT F97 variants and identifying beneficial enzyme mutants from a library of ATF1_Sc_, generated by model-guided protein design, for enhanced butyl acetate production. We anticipate the high-throughput microbial screening platform is a useful tool to identify novel AATs that have important roles in nature and industrial biocatalysis for designer bioester production.

## 2. MATERIALS AND METHODS

### 2.1 Strains and plasmids

Strains and plasmids used in this study are listed in Table 1. *E. coli* TOP10 was used for molecular cloning while BL21 (DE3) or EcDL002 (Layton and Trinh 2014) was used as a host strain for ester production. The pETDuet-1 plasmids containing 20 F97 variants of CAT_Sa_ were used to examine the role of the F97 residue on the alcohol substrate preference. The plasmid pATF1_Sc_ was constructed by subcloning *ATF1*_*Sc*_ gene from pDL004 (Layton and Trinh 2016a) into pET29 by the Gibson gene assembly method (Gibson et al. 2009). The ATF1_Sc_ variants were generated by site-directed mutagenesis (Zheng et al. 2004). All the constructed plasmids were introduced into the host strains by chemical transformation. The primers used in this study are listed in Table S1. The alphabets annotate the amino acid variants, including: R, arginine; H, histidine; K, lysine; D, aspartic acid; E, glutamic acid; S, serine; T, threonine; N, asparagine; Q, glutamine; C, cysteine; G, glycine; Y, tyrosine; P, proline; A, alanine; V, valine; I, isoleucine; L, leucine; M, methionine; F, phenylalanine; and W, tryptophan.

**Table 1:** The list of plasmids and strains used in this study.

| Name | Description | Source |
| --- | --- | --- |
| **Strains** |  |  |
| *E. coli* TOP10 | F<sup>-</sup> *mcrA* Δ(*mrr-hsdRMS-mcrBC*) φ80*lacZ*Δ*M15*<br>Δ*lacX74 recA1 araD139* Δ(*ara-leu*)7697 *galU galK*<br>λ<sup>-</sup> *rpsL*(Str<sup>R</sup>) *endA1 nupG* | Invitrogen |
| BL21 (DE3) | F<sup>-</sup> *ompT hsdS<sub>B</sub>* (r<sub>B</sub><sup>-</sup> m<sub>B</sub><sup>-</sup>) *gal dcm* (DE3) | Invitrogen |
| EcDL002 | TCS083 Δ*fadE* (DE3) | (Layton and Trinh 2014) |
| **Plasmids** |  |  |
| pCAT<sub>Sa</sub> | pETDuet-1 carrying CAT<sub>Sa</sub>; Amp<sup>R</sup> | (Seo et al. 2019) |
| pCAT<sub>Sa</sub> F97R | pETDuet-1 carrying CAT<sub>Sa</sub> F97R; Amp<sup>R</sup> | (Seo et al. 2019) |
| pCAT<sub>Sa</sub> F97H | pETDuet-1 carrying CAT<sub>Sa</sub> F97H; Amp<sup>R</sup> | (Seo et al. 2019) |
| pCAT<sub>Sa</sub> F97K | pETDuet-1 carrying CAT<sub>Sa</sub> F97K; Amp<sup>R</sup> | (Seo et al. 2019) |
| pCAT<sub>Sa</sub> F97D | pETDuet-1 carrying CAT<sub>Sa</sub> F97D; Amp<sup>R</sup> | (Seo et al. 2019) |
| pCAT<sub>Sa</sub> F97E | pETDuet-1 carrying CAT<sub>Sa</sub> F97E; Amp<sup>R</sup> | (Seo et al. 2019) |
| pCAT<sub>Sa</sub> F97S | pETDuet-1 carrying CAT<sub>Sa</sub> F97S; Amp<sup>R</sup> | (Seo et al. 2019) |
| pCAT<sub>Sa</sub> F97T | pETDuet-1 carrying CAT<sub>Sa</sub> F97T; Amp<sup>R</sup> | (Seo et al. 2019) |
| pCAT<sub>Sa</sub> F97N | pETDuet-1 carrying CAT<sub>Sa</sub> F97N; Amp<sup>R</sup> | (Seo et al. 2019) |
| pCAT<sub>Sa</sub> F97Q | pETDuet-1 carrying CAT<sub>Sa</sub> F97Q; Amp<sup>R</sup> | (Seo et al. 2019) |
| pCAT<sub>Sa</sub> F97C | pETDuet-1 carrying CAT<sub>Sa</sub> F97C; Amp<sup>R</sup> | (Seo et al. 2019) |
| pCAT<sub>Sa</sub> F97G | pETDuet-1 carrying CAT<sub>Sa</sub> F97G; Amp<sup>R</sup> | (Seo et al. 2019) |
| pCAT<sub>Sa</sub> F97Y | pETDuet-1 carrying CAT<sub>Sa</sub> F97Y; Amp<sup>R</sup> | (Seo et al. 2019) |
| pCAT<sub>Sa</sub> F97P | pETDuet-1 carrying CAT<sub>Sa</sub> F97P; Amp<sup>R</sup> | (Seo et al. 2019) |
| pCAT<sub>Sa</sub> F97A | pETDuet-1 carrying CAT<sub>Sa</sub> F97A; Amp<sup>R</sup> | (Seo et al. 2019) |
| pCAT<sub>Sa</sub> F97V | pETDuet-1 carrying CAT<sub>Sa</sub> F97V; Amp<sup>R</sup> | (Seo et al. 2019) |
| pCAT<sub>Sa</sub> F97I | pETDuet-1 carrying CAT<sub>Sa</sub> F97I; Amp<sup>R</sup> | (Seo et al. 2019) |
| pCAT<sub>Sa</sub> F97L | pETDuet-1 carrying CAT<sub>Sa</sub> F97L; Amp<sup>R</sup> | (Seo et al. 2019) |
| pCAT<sub>Sa</sub> F97M | pETDuet-1 carrying CAT<sub>Sa</sub> F97M; Amp<sup>R</sup> | (Seo et al. 2019) |
| pCAT<sub>Sa</sub> F97W | pETDuet-1 carrying CAT<sub>Sa</sub> F97W; Amp<sup>R</sup> | (Seo et al. 2019) |
| pDL004 | pETite carrying ATF1<sub>Sc</sub>; Kan<sup>R</sup> | (Layton and Trinh 2016a) |
| pATF1<sub>Sc</sub> | pET29 carrying ATF1<sub>Sc</sub>; Kan<sup>R</sup> | This study |
| pATF1<sub>Sc</sub> P348W | pET29 carrying ATF1<sub>Sc</sub> P348W; Kan<sup>R</sup> | This study |
| pATF1<sub>Sc</sub> P348R | pET29 carrying ATF1<sub>Sc</sub> P348R; Kan<sup>R</sup> | This study |
| pATF1<sub>Sc</sub> P348M | pET29 carrying ATF1<sub>Sc</sub> P348M; Kan<sup>R</sup> | This study |
| pATF1<sub>Sc</sub> P348H | pET29 carrying ATF1<sub>Sc</sub> P348H; Kan<sup>R</sup> | This study |
| pATF1<sub>Sc</sub> P348K | pET29 carrying ATF1<sub>Sc</sub> P348K; Kan<sup>R</sup> | This study |
| pATF1<sub>Sc</sub> P348N | pET29 carrying ATF1<sub>Sc</sub> P348N; Kan<sup>R</sup> | This study |
| pATF1<sub>Sc</sub> P348I | pET29 carrying ATF1<sub>Sc</sub> P348I; Kan<sup>R</sup> | This study |
| pATF1<sub>Sc</sub> P348S | pET29 carrying ATF1<sub>Sc</sub> P348S; Kan<sup>R</sup> | This study |
| pATF1<sub>Sc</sub> P348D | pET29 carrying ATF1<sub>Sc</sub> P348D; Kan<sup>R</sup> | This study |
| pATF1<sub>Sc</sub> P348C | pET29 carrying ATF1<sub>Sc</sub> P348C; Kan<sup>R</sup> | This study |
| pATF1<sub>Sc</sub> P348A | pET29 carrying ATF1<sub>Sc</sub> P348A; Kan<sup>R</sup> | This study |
| pATF1<sub>Sc</sub> P348Q | pET29 carrying ATF1<sub>Sc</sub> P348Q; Kan<sup>R</sup> | This study |

### 2.2 Culture media

The lysogeny broth (LB) medium, comprising of 10 g/L peptone, 5 g/L yeast extract, and 5 g/L NaCl, was used for molecular cloning and seed cultures. The M9 hybrid medium (Layton and Trinh 2014) with 20 g/L glucose was used for ester production. Either 50 µg/mL ampicillin (Amp) or 50 µg/mL kanamycin (Kan) was added to the media for selection where applicable.

### 2.3 Microplate-based microbial screening method

Cell inoculum was prepared either from a bacterial glycerol stock or from a single colony on a LB agar plate. Specifically, 1% (v/v) of stock cells were grown overnight in 5 mL of LB at 37°C and 200 rpm on a 75° angled platform in a New Brunswick Excella E25 (Eppendorf, CT, USA). Alternatively, single colonies from LB agar plates were inoculated in 100 µL of LB in 96-well microplates using sterile pipette tips. Each colony picked by a sterile pipette tip was subsequently mixed with the media in the target well and was grown overnight at 37°C and 400 rpm in an incubating microplate shaker.

For the microplate-based screening assay, 5 % (v/v) of cell inocula were first inoculated in 100 µL of the M9 hybrid media containing 20 g/L of glucose, 0.1 mM of isopropyl β-D-1-thiogalactopyranoside (IPTG), and 2 g/L of alcohol (i.e., ethanol, n-butanol, isobutanol, or 2-phenylethyl alcohol) in 96-well microplates with or without hexadecane overlay in a 1:1 (v/v) ratio. The microplates were then sealed with a plastic adhesive sealing film, SealPlate® (EXCEL Scientific, Inc., CA, USA), to avoid cross contamination and evaporation. Finally, the microplates were incubated at 37°C and 400 rpm for 24 hours (h) in an incubating microplate shaker (Fisher Scientific, PA, USA).

The optical density (OD) of cell culture was measured at 600 nm using a BioTek Synergy HT microplate reader (BioTek Instruments, Inc., VT, USA). The dry cell weight (DCW) was obtained by multiplication of the optical density of culture broth with a pre-determined conversion factor, 0.385 g/L/OD. The organic layers were collected for ester measurement either by gas chromatography coupled with mass spectroscopy (GC/MS) or colorimetric assay.

### 2.4 SDS-PAGE analysis

For SDS-PAGE analysis, 1% (v/v) of stock cells were grown overnight at 37°C and 200 rpm in 15 mL culture tubes containing 5 mL of LB media and antibiotics. Then, 4% (v/v) of the overnight cultures were transferred into 1.5 mL of LB media containing antibiotics in a 24-well microplate (cat# 353224, BD Falcon). The cultures were next grown at 37°C and 400 rpm using an incubating microplate shaker. When the cultures reached an OD of 0.4∼0.6, they were induced with 0.1 mM IPTG and sealed with a Breathe-Easy Sealing Membrane to prevent evaporation and cross contamination (cat# BEM-1, Research Products International Corp., IL, USA). After 4 h of induction, cells were collected by centrifugation and resuspended in 1X phosphate-buffered saline (PBS) buffer (pH7.4) at the final OD of 5. The cell pellets were disrupted using the B-PER complete reagent (cat# 89822, Thermo Scientific, MA, USA). The resulting crude extracts were mixed with 6x sodium dodecyl sulfate (SDS) sample buffer and heated at 95°C for 5 min. Finally, the protein samples were analyzed by SDS-polyacrylamide gel electrophoresis (SDS-PAGE, 14% polyacrylamide gel). Protein bands were visualized with Coomassie Brilliant Blue staining.

### 2.5 Gas chromatography coupled with mass spectroscopy (GC/MS)

The microplates from the microbial screening of AATs were centrifuged at 4,800 x g for 5 min and the hexadecane overlays were used for quantification of esters. The samples were prepared by diluting hexadecane extracts from the cultures with hexadecane containing internal standard (isoamyl alcohol) in a 1:1 (v/v) ratio. Then, 1 μL of samples were directly injected into a gas chromatograph (GC) HP 6890 equipped with the mass selective detector (MS) HP 5973. For the GC system, helium was used as the carrier gas at a flow rate of 0.5 mL/min and the analytes were separated on a Phenomenex ZB-5 capillary column (30 m x 0.25 mm x 0.25 μm). The oven temperature was programmed with an initial temperature of 50°C with a 1°C/min ramp to 58°C. Next a 25°C/min ramp was deployed to 235°C and then a 50°C/min ramp was deployed to 300°C. Finally held a temperature of 300°C for 2 minutes to elute any residual non-desired analytes. The injection was performed using the splitless mode with an initial injector temperature of 280°C. For the MS system, a selected ion monitoring (SIM) mode was deployed to detect analytes. The SIM parameters for detecting esters were as follows: i) for ethyl acetate, ions 45.00, and 61.00 detected from 4.15 to 5.70 min, ii) for isoamyl alcohol (internal standard), ions 45.00, and 88.00 detected from 5.70 to 6.60 min, iii) for isobutyl acetate, ions 61.00, and 101.00 detected from 6.60 to 7.75 min, iv) for butyl acetate, ions 61.00, and 116.00 detected from 7.75 to 13.70 min, and v) for 2-phenethyl acetate, ions 104.00, and 121.00 detected from 13.70 to 13.95 min.

### 2.6 Colorimetric assay for ester quantification

The colorimetric assay for ester quantification was performed in a 96-well microplate. The protocol was slightly modified from the previously reported method (Lobs et al. 2016). Specifically, in each well, 60 µL of hexadecane overlay from the culture was mixed with 20 µL of hydroxylamine stock solution and incubated in an incubating microplate shaker at room temperature for 10 minutes (min) to produce hydroxamic acid. Next, 120 µL of the ferric working solution (1/20-diluted stock ferric iron(III) solution in ethanol) was added to the reaction solution and incubated for 5 min to form an iron-hydroxamic acid complex. Finally, the absorbance was measured at 520 nm (Ab_520_) using a BioTek Synergy HT microplate reader. Esters were quantified using a standard curve between the absorbances and known concentrations of a target ester.

### 2.7 *In silico* mutagenesis of AATs

The 3D structures of ATF1_Sc_, acetyl-CoA, and butanol were first prepared for the model-guided protein engineering of ATF1_Sc_ in MOE (Molecular Operating Environment software), as previously described (Seo et al. 2019). To perform docking simulations in MOE, the potential binding pocket of ATF1_Sc_ was identified using the ‘Site Finder’ tool. Both of the conserved catalytic residues, H191 and D195, that are known to reside in the binding pocket of AATs (Navarro-Retamal et al. 2016), were selected. Acetyl-CoA and butanol were docked to the binding pocket of ATF1_Sc_ in a sequential step. After the docking simulations, the best-scored binding poses were selected for *in silico* mutagenesis. The ‘residue scan’ tool of MOE was used to identify beneficial mutations for the biosynthesis of the target ester (i.e., butyl acetate) in ATF1_Sc_ variants, based on their ΔAffinity (kcal/mol) values. Here, the ΔAffinity value represents the relative binding affinity between a mutant and its wild type, where a more negative value indicates a mutant with higher affinity. Details of the docking simulation and *in silico* mutagenesis analysis protocols in MOE can be found in the previous report (Lee et al. 2018).

## 3. RESULTS AND DISCUSSION

### 3.1 Establishing a microplate-based culturing technique with *in situ* fermentation and extraction of esters

To develop a high-throughput microbial screening platform for ester biosynthesis, we first examined whether the microplate-based culturing technique could be reliably used to monitor cell growth and continuously extract esters for downstream quantification. We characterized the recombinant BL21 (DE3) strains harboring 20 CAT_Sa_ F97 variants (Seo et al. 2019) in 96-well microplates for IBA production with or without solvent (hexadecane) overlay for *in situ* ester extraction (Fig. 1A). As a basis for comparison, we also performed the high cell density culturing method in shake tubes that was previously employed for AAT screening (Seo et al. 2019).

**Figure 1.**
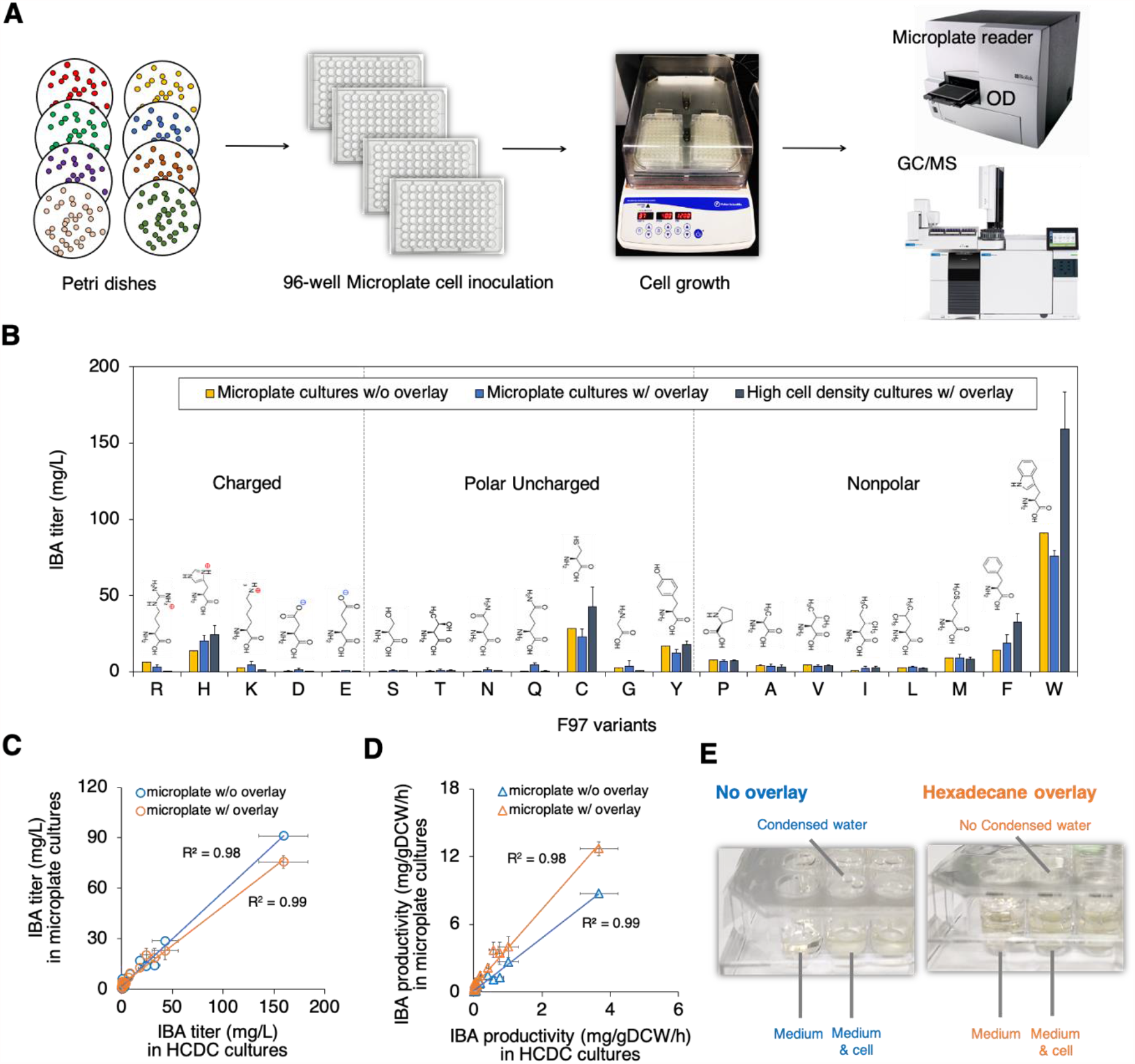
A microplate-based culturing method for microbial biosynthesis of esters with *in situ* product extraction. **(A)** Workflow of the microplate-based culturing method. (**B**) Comparison of IBA production among three different culturing methods by the recombinant BL21 (DE3) strains carrying the 20 CAT_Sa_ F97 variants after 24 h. (**C-D**) Comparison of **(C)** IBA titer and **(D)** specific productivity among three different culturing methods. The error bars represent standard deviation of four biological replicates (*n*=4). (**E**) Effect of solvent overlay on growth kinetics measurement in the microplate-based culturing method.

The characterization results show that IBA production in microplates followed the same trend of its production observed in high cell density cultures (Fig. 1B). Strong positive linear correlations (R^2^ ≥ 0.98) in IBA production existed between the microplate-based and high cell density culturing methods (Figs. 1C, 1D). The microplate-based culturing method could validate that the CAT_Sa_ F97W variant achieved the highest IBA production among the 20 characterized variants (Table S2). While these variants exhibited different activities toward IBA, their cell cultures exhibited similar growth (Fig. S1).

Using solvent overlays for microbial biosynthesis of esters can provide several key advantages. First, the solvent overlay enables a more reliable measurement of cell growth in microplates by avoiding medium evaporation that caused water condensation (Fig. 1E) and hence interfered with optical density measurement (Fig. S1). Growth kinetics helps evaluate the effect of expressing AATs on microbial health. Second, the solvent overlay simplifies the sample preparation step for quantification of esters and is compatible with a high-throughput workflow, where esters in the solvent layer can be extracted for downstream measurement by either a GC/MS method or a colorimetric assay (Layton and Trinh 2014; Layton and Trinh 2016a; Layton and Trinh 2016b; Lobs et al. 2016; Rodriguez et al. 2014). Third, the solvent overlay helps alleviate the product toxicity during fermentation (Brennan et al. 2012) because esters are known to be inhibitory to microbial health (Wilbanks and Trinh 2017).

Taken altogether, the microplate-based culturing method with solvent overlays is reliable and suitable for a rapid, high-throughput *in vivo* screening platform for microbial ester production.

### 3.2 Revealing the influential role of CAT_Sa_ F97 residue by profiling alcohol preference of a library of F97 mutants with the microplate-based culturing method

Our previous study discovered that the CAT_Sa_ F97W mutant improved its catalytic efficiency towards isobutanol by ∼2-fold (Seo et al. 2019). We hypothesized that the F97 residue might have an important role in determining the alcohol substrate preference. Using the established microplate-based culturing method, we evaluated whether it could be used for rapid profiling of the alcohol substrate preference of CAT_Sa_ F97 variants. We characterized the recombinant *E. coli* strains carrying 20 CAT_Sa_ F97 variants with exogenous supplementation of alcohols in the media including linear, short-chain alcohols (ethanol, butanol), a branched-chain alcohol (isobutanol), and an aromatic alcohol (2-phenylethyl alcohol) in microplates with hexadecane overlay.

The characterization results showed that mutations in the F97 residue changed the ester production profiles (Fig. 2A, Table S3), suggesting that F97 plays an important role in determining the alcohol substrate preference of CAT_Sa_. All the CAT_Sa_ variants exhibited poor activities toward ethanol (Table S3). Among the four target acetate esters investigated, F97H produced 2-phenylethyl acetate (PEA) at the highest level of 194.63 mg/L followed by F97H (182.30 mg/L) and the wildtype F97 (149.41 mg/L) (Fig. 2A, Table S3). As compared to the wildtype, IBA production by F97W (91.22 mg/L) showed the highest improvement (∼6.39-fold) (Fig. 2A, Table S3), which was relatively consistent with the prior *in vitro* study showing that F97W variant achieved ∼2-fold increase in the catalytic efficiency towards isobutanol (Seo et al. 2019). Remarkably, F97T showed BA production (12.42 mg/L) with high specificity, demonstrating the feasibility of production of designer esters using re-programmed CATs (Fig. 2A, Table S3). Different from F97W, F97C exhibited the highest n-butyl acetate (BA) production (18.26 mg/L) (Fig. 2A, Table S3). Examining the protein structure of CAT_Sa_ can provide some insights its alcohol substrate preference (Fig. 2B). As shown, the binding pockets of CATs are formed at the subunit interfaces (Day and Shaw 1992) and the F97 residue located on the opposite subunit of the catalytic residues, H189 and D193. One possible explanation on how only one residue replacement influences the substrate preference of CAT_Sa_ is that the mutation in the F97 residue might dramatically change the size and/or shape of the binding pocket and hence alternate the interactions among subunits of CAT_Sa_.

**Figure 2.**
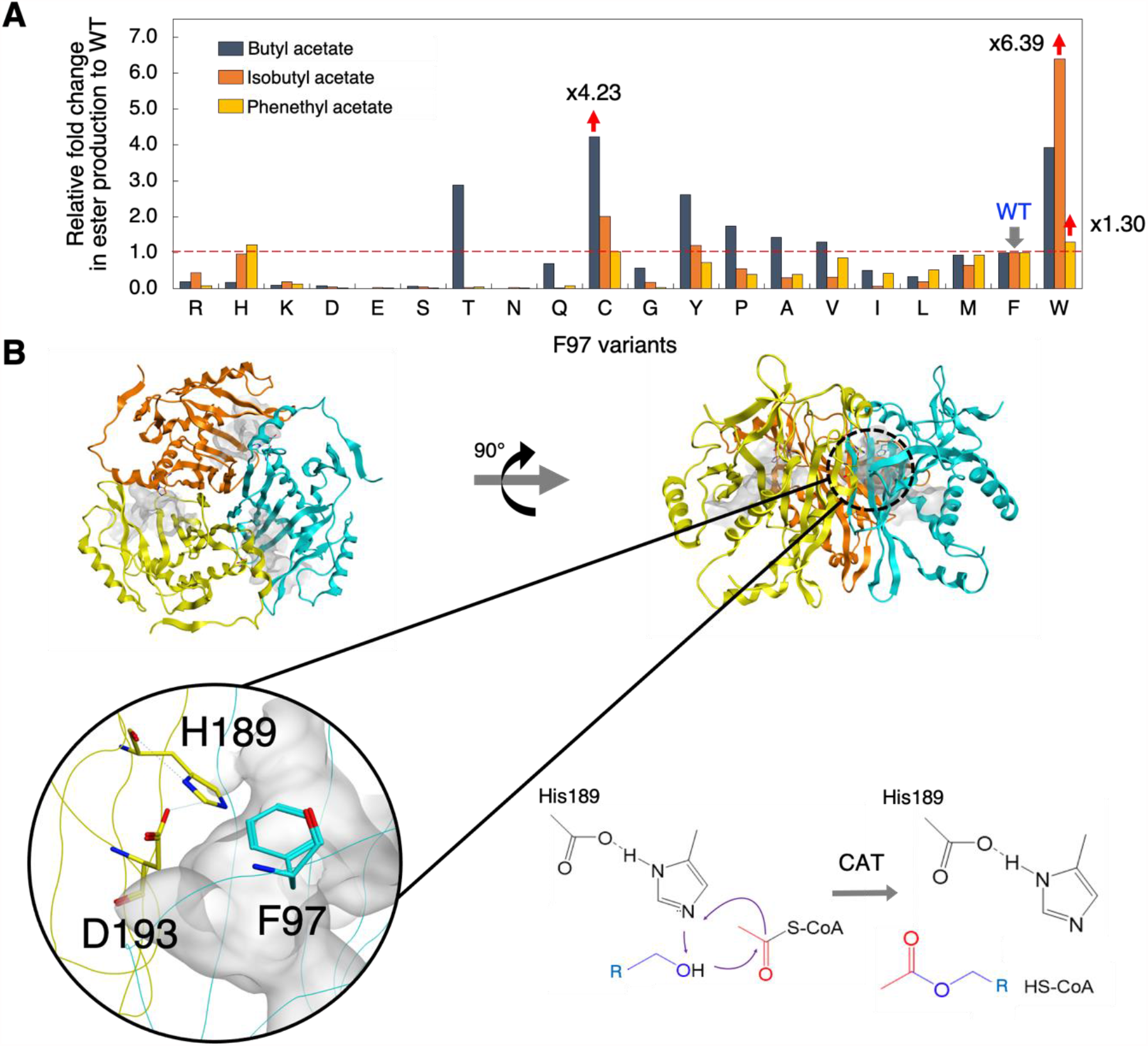
Rapid profiling of the alcohol substrate preference of CAT_Sa_ F97 variants. (**A**) Relative fold change in ester production to wild type. (**B**) The 3D structure of homology model of CAT_Sa_ and the reaction mechanism. F97 residue (in green); catalytic residues H189 and D193 (in cyan); binding pockets (in grey clouds). Note: ethyl acetate production was not shown here due to low or no detectable amounts (Table S3).

Taken altogether, the microplate-based culturing method coupled with GC/MS can be employed for rapid profiling of substrate preferences of AATs. The method revealed the important role of the F97 residue in determining the alcohol substrate preference of CAT_Sa_.

### 3.3 Developing a high-throughput microbial screening platform for ester biosynthesis by integration of the microplate-based culturing method and a colorimetric assay

High-throughput screening of ester biosynthesis has been limited by the use of GC/MS. Since the reactions of esters with hydroxylamine generate hydroxamic acids that form purple complexes with ferric ion, esters can be determined colorimetrically by measuring absorbance at 520 nm (Fig. 3A, 3B) (Hill 1946; Lobs et al. 2016; Stern and Shapiro 1953; Wofford et al. 1986). This colorimetric assay has recently been adapted for high-throughput screening of ethyl acetate (EA) production from C5, C6, and C12 carbon sources in *Kluyveromyces marxianus* (Lobs et al. 2016) where cell culture samples were first collected followed by ester extraction with hexane. This protocol is useful but might not be compatible with the microplate-based culturing method in our study because hexane is toxic and hence cannot be used for *in situ* fermentation and extraction, unlike hexadecane. Here, we tested whether the colorimetric assay can be modified and coupled with the microplate-based culturing method to facilitate a high-throughput microbial screening of AATs for ester biosynthesis.

**Figure 3.**
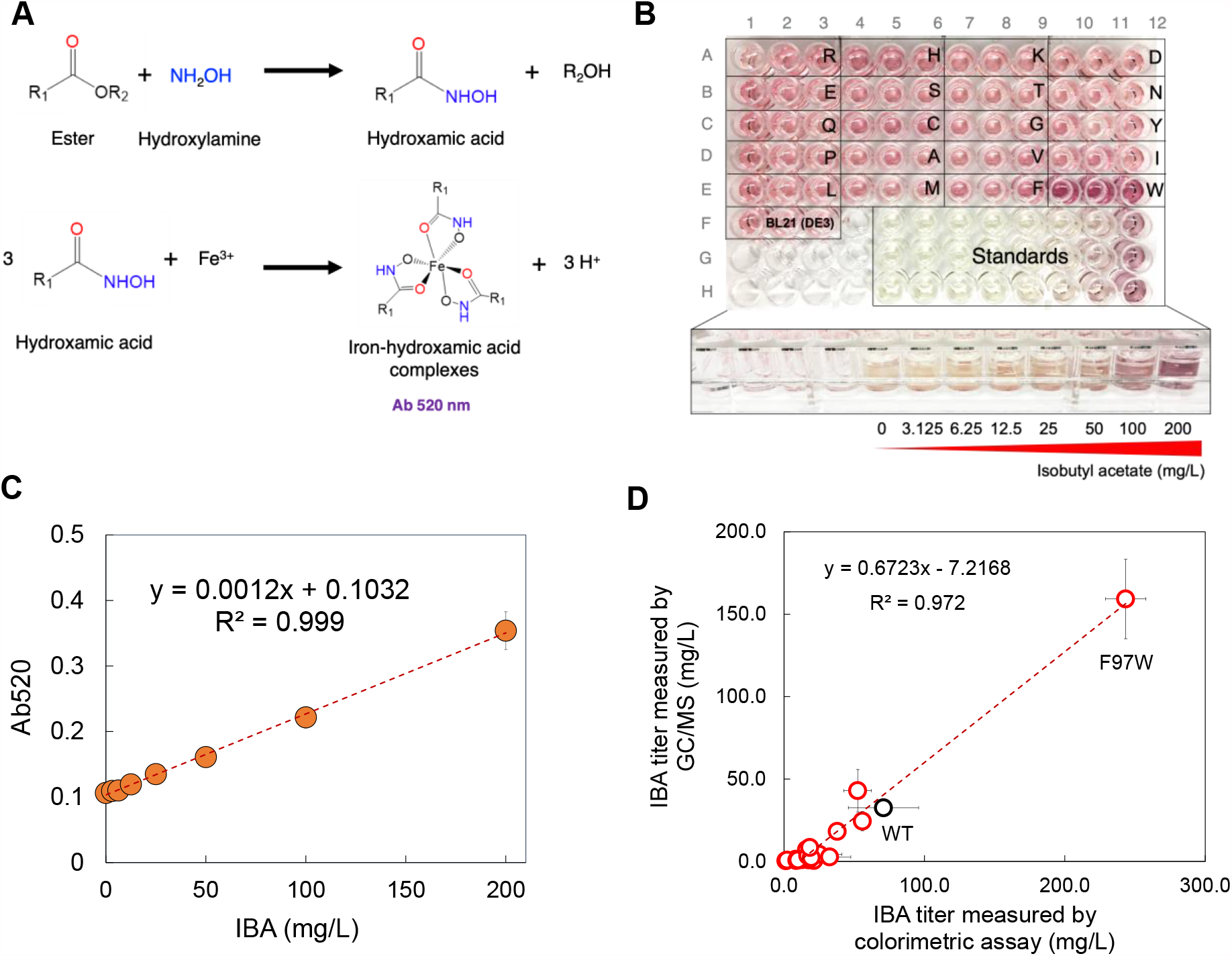
Development of the high-throughput microbial screening platform. (**A**) Chemical reactions involved in the colorimetric assay for ester quantification. (**B**) Demonstration of the colorimetric assay in a microplate conducted for hexadecane overlay samples and a series of IBA standards at different concentrations. (**C**) The IBA standard curve. (**D**) Comparison of the IBA titers measured by the high-throughput microbial screening and GC/MS methods. The error bars represent standard deviation of three biological replicates (*n*=3).

To compare ester quantification by the colorimetric assay and GC/MS method, we analyzed IBA production by the *E. coli* BL21 (DE3) strains harboring the 20 F97 variants as demonstrated for microplate-based culturing method. We started by developing a standard curve to estimate IBA production by the colorimetric assay. Using pure IBA in hexadecane, an almost perfect linear correlation (R^2^=0.999) was established between absorbance at 520 nm and IBA concentration within 0-200 mg/L (Fig. 3C). When using esters in hexadecane from the cell culture samples for the colorimetric assay, we found that the colorimetric assay could determine the IBA concentrations consistently with the GC/MS method (Fig. 3D).

Critical to the high-throughput microbial screening method to estimate the target esters from the culture samples is to have an appropriate control for the baseline adjustment in the colorimetric assay. In our study, we found that the colorimetric method could overestimate the IBA production as compared to the GC/MS method (Fig. S2B, Table S4). Since the *E. coli* host produced ethanol endogenously, EA was produced as an inevitable by-product (Fig. S2A), causing the observed IBA overestimation. To avoid this problem, we used ΔAb_520_ for estimating IBA production where ΔAb_520_ = Ab_520, AAT_^+^,_ROH_^+^ - Ab_520, AAT_^+^,_ROH_^-^ is the absorbance difference between culture samples with and without the target alcohol (ROH) availability. The target alcohol can be supplemented externally or produced by the cells. One other strategy that might help avoid the target ester (e.g., IBA) overestimation problem by the colorimetric assay is to use a host strain void of the endogenous pathways causing the biosynthesis of the unwanted alcohol byproduct (e.g., ethanol).

It is important to note that during our protocol development, we observed that the perturbed, emulsified layer of an immiscible hexadecane-ethanol mixture interfered with the measured absorbance and generated irreproducible data. Note that ethanol is originated from the ferric solution used in the colorimetric assay. This problem did not occur in the previous study (Lobs et al. 2016) likely because hexane used for ester extraction is miscible in ethanol. To address this problem, we used a centrifugation step to create the immiscible hexadecane-ethanol mixture with the transparent organic phase and strong purple aqueous phase (Fig. 3B).

Overall, the microplate-based culturing method coupled with a colorimetric assay is suitable for high-throughput microbial screening of AATs for ester biosynthesis.

### 3.4 Combining the model-guided protein engineering and high-throughput microbial screening platform to rapidly identify ATF1_Sc_ variants for improved BA production

With the established high-throughput microbial screening platform, we applied it to rapidly identify the engineered ATF1_Sc_ mutants for enhanced BA production (Fig. 4A). We started by generating a library of potential ATF1_Sc_ candidates for improved BA production *in silico* for high-throughput microbial screening. To do this, we first created a 3-D structure of ATF1_Sc_ using the homology model of 15-O-acetyltransferase (PDB:3FP0) best predicted by SWISS-MODEL (Waterhouse et al. 2018). We next identified the binding pocket of ATF1_Sc_ for docking simulations of the BA co-substrates, including acetyl-CoA and butanol. Based on the homology model, the binding pocket of ATF1_Sc_ consists of 24 residues including V32, Y36, H191, D195, G196, R197, T316, I347, P348, A349, D350, R352, N370, V371, I374, F376, Y399, I403, L407, K426, L448, S449, N450, V451, F471, and Q473, where H191 and D195 are the catalytic residues (Fig. S3A). By performing docking simulations, we generated the acetyl-CoA-butanol-ATF1_Sc_ complex and identified the residues interacting with butanol including V32, Y36, D195, P348, V371, L447, S449, Q473, Q475, and S483 (Fig. S3B). Finally, we performed the residue scan against these 10 residues to select the top 12 promising candidates including P348W, P348R, P348M, P348H, P348K, P348N, P348I, P348S, P348D, P348C, P348A, and P348Q for experimental characterization (Fig. 4B).

**Figure 4.**
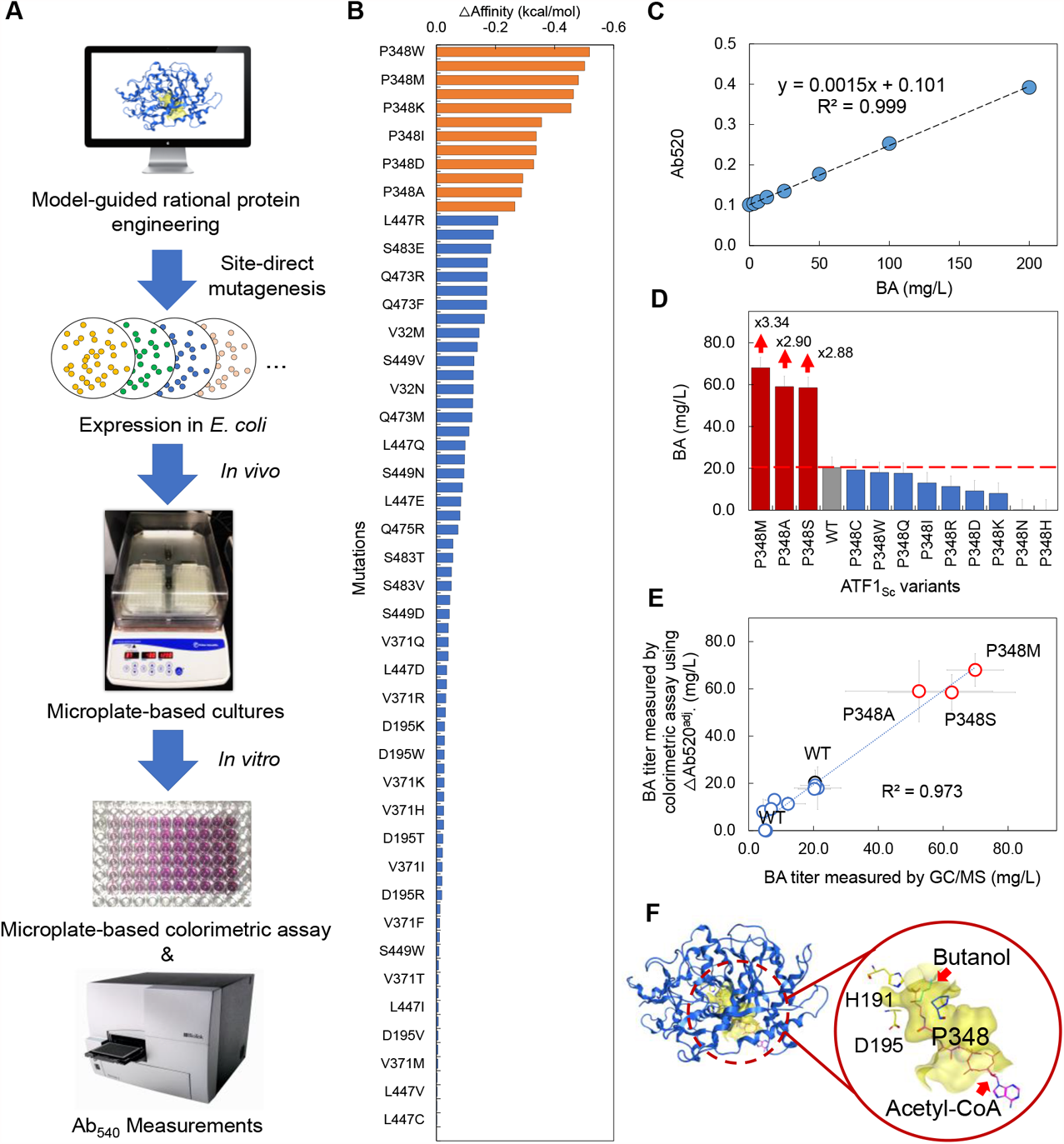
Rapid identification of the beneficial ATF1_Sc_ variants for improved butyl acetate (BA) production. (**A**) A schematic workflow of the high-throughput microbial screening platform used to identify ATF1_Sc_ from a library of variants generated by the model-aided protein design. (**B**) Residue scan results of the residues interacting with butanol in the acetyl-CoA-butanol-ATF1_Sc_ complex. The orange bars represent the selected top 12 candidates for further studies. (**C**) The BA standard curve used for the quantification of BA in the colorimetric assay. (**D**) Screening results of the selected top 12 candidates. The error bars represent standard deviation of four biological replicates (*n*=4). (**E**) Correlation of the measured BA titer between the colorimetric assay and GC/MS methods. (**F**) The location of P348 residue in ATF1_Sc_. A yellow cloud represents a binding pocket of ATF1_Sc_.

To perform the high-throughput microbial screening of the top 12 engineered ATF1_Sc_ candidates for improved BA production, we used TCS083 Δ*fadE* (DE3) (Layton and Trinh 2014) as a host strain. Like the colorimetric assay developed for IBA measurement, the standard curve for BA measurement showed a strong linear correlation (R^2^=0.999) between the 520 nm absorbance and the standard BA concentrations in the range of 0-200 mg/L (Fig. 4C). Our screening results shows that the P348M, P348A, or P348S mutation in ATF1_Sc_ improved BA production by 3.34, 2.90, or 2.88-fold, respectively (Fig. 4D). To confirm this result, we compared the BA titers measured by the colorimetric assay with those by GC/MS. Remarkably, we could observe almost identical BA titers between the two methods (Table S5) with a strong linear correlation (R^2^=0.973) (Fig. 4E). This result demonstrates the high-throughput microbial screening platform is suitable for rapidly identifying AATs for designer ester biosynthesis. Interestingly, like the F97 residue in CAT_Sa_ (Fig. 2B), the P348 residue in ATF1_Sa_ is also located on the opposite side of the catalytic residues including H191 and D195 (Fig. 4F) and interacts with an alcohol substrate, which might determine the alcohol substrate preference.

Overall, we rationally engineered ATF1_Sc_ for improved BA production through a model-guided rational protein engineering and rapidly identified the beneficial ATF1_Sc_ variants using the established high-throughput microbial screening platform.

## 4. CONCLUSION

We developed a high-throughput microbial screening platform to probe specificities of AATs/CATs for designer ester biosynthesis. This platform integrated the microplate culturing method with a modified colorimetric assay previously established, which provides useful information about AAT expression and activity, microbial health, and ester production. For the microplate-based culturing protocol, the use of solvent overlays is critical to minimize medium evaporation, generate reproducible growth measurement, and eliminate the ester extraction step. For colorimetric assay, the addition of a centrifugation step is crucial to avoid the interference of ethanol-hexadecane immiscible layer that causes irreproducible measurement. The high-throughput microbial screening platform not only confirmed CAT_Sa_ F97W with enhanced isobutyl acetate synthesis but also identified the three ATF1_Sc_ (P348M, P348A, and P348S) variants generated by model-guided rational protein engineering for enhanced butyl acetate production. Overall, this study presents a high-throughput microbial screening platform for rapid profiling of the alcohol substrate preference of AATs for production of designer esters. We believe that this platform is scalable and compatible with automated microplate handling systems to increase its screening capacity.

## ACKNOWLEDGEMENTS

This research was financially supported in part by the NSF CAREER award (NSF#1553250), the DOE BER Genomic Science Program (DE-SC0019412), and the DOE subcontract grant (DE-AC05-000R22725) by the Center of Bioenergy Innovation, the U.S. Department of Energy Bioenergy Research Center funded by the Office of Biological and Environmental Research in the DOE Office of Science, and the U.S. Department of Energy Joint Genome Institute. The authors would like to thank the Center of Environmental Biotechnology at UTK for using the GC/MS instrument.

