## Supplementary Materials for "Probing Specificities of Alcohol Acyltransferases for Designer Ester Biosynthesis with a High-Throughput Microbial Screening Platform"

**Supplementary Table S1.** List of primers used in this study. The underlined letters indicate site-directed mutation sites of ATF1$_{Sc}$. F, R, and BB represent forward, reverse, and backbone, respectively.

| Primers | Primer sequence (5' to 3') | Description |
|---|---|---|
| ATF1$_{Sc}$_F | ATAATTTTGTTTAACTTTAAGAAGGAGATATAGATA TGAATGAAATCGATGAGAAAAAT | Primers for constructing the pATF1$_{Sc}$ by Gibson assembly |
| ATF1$_{Sc}$_R | TTTGTTAGCAGCCGGATCTCAGTGGTGGTGGTGGTG GTGGATAGGGCCTAAAAGGAGAG | |
| BB_ATF1$_{Sc}$_F | ATCCACCACCACCACC | |
| BB_ATF1$_{Sc}$_R | ATCTATATCTCCTTCTTAAAGTTAAACAAAATTATT TCTAG | |
| P348W F | ATTTTTATCTGGGCAGATTGCCGCTCACAACTA | Primers for constructing the pATF1$_{Sc}$ variants by site-directed mutagenesis using the QuickChange™ site-directed mutagenesis kit |
| P348W R | GGCAATCTGCCCAGATAAAAATATCCGTAAGCC | |
| P348R_F | ATTTTTATCCGTGCAGATTGCCGCTCACAACTA | |
| P348R_R | GGCAATCTGCACGGATAAAAATATCCGTAAGCC | |
| P348M_F | ATTTTTATCATGGCAGATTGCCGCTCACAACTA | |
| P348M_R | GGCAATCTGCCATGATAAAAATATCCGTAAGCC | |
| P348H_F | ATTTTTATCCATGCAGATTGCCGCTCACAACTA | |
| P348H_R | GGCAATCTGCATGGATAAAAATATCCGTAAGCC | |
| P348K_F | ATTTTTATCAAAGCAGATTGCCGCTCACAACTA | |
| P348K_R | GGCAATCTGCTTTGATAAAAATATCCGTAAGCC | |
| P348N_F | ATTTTTATCAATGCAGATTGCCGCTCACAACTA | |
| P348N_R | GGCAATCTGCATTGATAAAAATATCCGTAAGCC | |
| P348I_F | ATTTTTATCATTGCAGATTGCCGCTCACAACTA | |
| P348I_R | GGCAATCTGCAATGATAAAAATATCCGTAAGCC | |
| P348S_F | ATTTTTATCACAGCAGATTGCCGCTCACAACTA | |
| P348S_R | GGCAATCTGCTGTGATAAAAATATCCGTAAGCC | |
| P348D_F | ATTTTTATCGATGCAGATTGCCGCTCACAACTA | |
| P348D_R | GGCAATCTGCATCGATAAAAATATCCGTAAGCC | |
| P348C_F | ATTTTTATCTGCGCAGATTGCCGCTCACAACTA | |
| P348C_R | GGCAATCTGCGCAGATAAAAATATCCGTAAGCC | |
| P348A_F | ATTTTTATCGCTGCAGATTGCCGCTCACAACTA | |
| P348A_R | GGCAATCTGCAGCGATAAAAATATCCGTAAGCC | |
| P348Q_F | ATTTTTATCCAGGCAGATTGCCGCTCACAACTA | |
| P348Q_R | GGCAATCTGCCTGGATAAAAATATCCGTAAGCC | |

**Supplementary Table S2.** Summary of isobutyl acetate production after 24h by BL21 (DE3) strains carrying the 20 $CAT_{Sa}$ F97 variants using the various cell culturing methods. WT, wild type.

| F97 variants | Culture methods | | | | | |
|---|---|---|---|---|---|---|
| | Titer (mg/L) | | | Specific ester productivity (mg/gDCW/h) | | |
| | Microplates w/o overlay | Microplates w/ overlay | High cell density w/ overlay | Microplates w/o overlay | Microplates w/ overlay | High cell density w/ overlay |
| R | 6.28 ± 0.98 | 3.49 ± 1.02 | 0.57 ± 0.12 | 0.57 ± 0.00 | 0.59 ± 0.17 | 0.01 ± 0.00 |
| H | 13.67 ± 4.44 | 20.42 ± 3.78 | 24.43 ± 5.83 | 1.13 ± 0.01 | 3.75 ± 0.69 | 0.58 ± 0.14 |
| K | 2.63 ± 1.01 | 4.62 ± 2.21 | 1.29 ± 0.21 | 0.22 ± 0.01 | 0.75 ± 0.36 | 0.03 ± 0.00 |
| D | 0.62 ± 0.23 | 1.60 ± 0.53 | 0.37 ± 0.06 | 0.05 ± 0.01 | 0.26 ± 0.09 | 0.01 ± 0.00 |
| E | 0.43 ± 0.31 | 0.96 ± 0.23 | 0.55 ± 0.11 | 0.03 ± 0.00 | 0.16 ± 0.04 | 0.01 ± 0.00 |
| S | 0.68 ± 0.19 | 0.96 ± 0.29 | 0.90 ± 0.29 | 0.06 ± 0.00 | 0.17 ± 0.05 | 0.02 ± 0.01 |
| T | 0.59 ± 0.23 | 0.88 ± 1.02 | 1.04 ± 0.41 | 0.05 ± 0.02 | 0.16 ± 0.18 | 0.02 ± 0.01 |
| N | 0.42 ± 0.38 | 1.53 ± 1.24 | 0.78 ± 0.14 | 0.04 ± 0.01 | 0.26 ± 0.21 | 0.02 ± 0.00 |
| Q | 0.35 ± 0.56 | 4.56 ± 1.53 | 0.68 ± 0.10 | 0.04 ± 0.02 | 0.83 ± 0.28 | 0.02 ± 0.00 |
| C | 28.52 ± 5.22 | 22.96 ± 5.36 | 42.93 ± 12.84 | 2.70 ± 0.00 | 4.02 ± 0.94 | 1.01 ± 0.30 |
| G | 2.59 ± 1.65 | 3.90 ± 3.62 | 0.86 ± 0.18 | 0.25 ± 0.01 | 0.66 ± 0.61 | 0.02 ± 0.00 |
| Y | 17.08 ± 1.15 | 12.70 ± 2.26 | 18.21 ± 2.06 | 1.51 ± 0.01 | 2.15 ± 0.38 | 0.42 ± 0.05 |
| P | 7.86 ± 0.68 | 6.96 ± 0.71 | 7.20 ± 0.69 | 0.72 ± 0.01 | 1.06 ± 0.11 | 0.17 ± 0.02 |
| A | 4.29 ± 0.50 | 3.89 ± 1.18 | 3.44 ± 1.03 | 0.41 ± 0.01 | 0.60 ± 0.18 | 0.08 ± 0.02 |
| V | 4.60 ± 0.40 | 3.85 ± 0.72 | 4.30 ± 0.26 | 0.41 ± 0.01 | 0.66 ± 0.12 | 0.10 ± 0.01 |
| I | 1.04 ± 0.50 | 2.23 ± 1.34 | 2.70 ± 0.88 | 0.10 ± 0.01 | 0.38 ± 0.23 | 0.06 ± 0.02 |
| L | 2.66 ± 0.37 | 3.20 ± 0.54 | 2.32 ± 0.68 | 0.27 ± 0.01 | 0.53 ± 0.09 | 0.05 ± 0.02 |
| M | 9.17 ± 1.75 | 9.36 ± 2.21 | 8.44 ± 1.12 | 0.78 ± 0.01 | 1.47 ± 0.35 | 0.20 ± 0.03 |
| F (WT) | 14.17 ± 2.99 | 18.88 ± 5.52 | 32.55 ± 5.48 | 1.31 ± 0.01 | 3.46 ± 1.01 | 0.76 ± 0.13 |
| W | 90.53 ± 41.72 | 75.73 ± 3.70 | 159.26 ± 24.12 | 8.74 ± 0.01 | 12.72 ± 0.62 | 3.66 ± 0.55 |

**Supplementary Table S3.** The results of rapid profiling of the alcohol substrate preference of the 20 F97 $CAT_{Sa}$ variants by the microplate-based culturing method from the 24 h cultures. WT, wild type; *n.d.*, not detected.

| F97 variants | Ester titer (mg/L) | | | |
|---|---|---|---|---|
| | Ethyl acetate | Butyl acetate | Isobutyl acetate | 2-Phenylethyl acetate |
| R | *n.d.* | 0.83 | 6.33 | 11.82 |
| H | *n.d.* | 0.79 | 13.77 | 182.30 |
| K | *n.d.* | 0.44 | 2.65 | 19.74 |
| D | *n.d.* | 0.38 | 0.63 | 3.83 |
| E | *n.d.* | *n.d.* | 0.44 | 1.40 |
| S | *n.d.* | 0.31 | 0.69 | 3.52 |
| T | *n.d.* | 12.42 | 0.60 | 7.15 |
| N | *n.d.* | *n.d.* | 0.43 | 3.25 |
| Q | *n.d.* | 3.02 | 0.36 | 12.14 |
| C | *n.d.* | 18.26 | 28.74 | 153.91 |
| G | *n.d.* | 2.51 | 2.61 | 4.46 |
| Y | *n.d.* | 11.25 | 17.20 | 109.88 |
| P | *n.d.* | 7.53 | 7.92 | 58.66 |
| A | *n.d.* | 6.13 | 4.33 | 59.05 |
| V | *n.d.* | 5.64 | 4.63 | 126.99 |
| I | *n.d.* | 2.22 | 1.05 | 63.27 |
| L | *n.d.* | 1.47 | 2.68 | 79.27 |
| M | *n.d.* | 4.01 | 9.24 | 138.99 |
| F (WT) | *n.d.* | 4.32 | 14.28 | 149.41 |
| W | 0.39 | 16.93 | 91.22 | 194.63 |

**Supplementary Table S4.** Comparison of ester titers measured by the high-throughput microbial screening platform and GC/MS method. $\triangle Ab_{520} = Ab_{520,CAT^{+},IBOH^{+}} - Ab_{520,CAT^{+},IBOH^{-}}$. WT, wild type; *n.d.*, not detected. The used hexadecane overlay samples were obtained from high cell density cultures (2).

| $CAT_{Sa}$ F97 variants | GC/MS | | | Colorimetric assay |
|---|---|---|---|---|
| | Ester titer (mg/L) | | | $\triangle Ab_{520}$ |
| | Ethyl acetate | Isobutyl acetate | Total | |
| R | *n.d.* | 0.57 ± 0.12 | 0.57 ± 0.12 | 0.03 ± 0.00 |
| H | 0.03 ± 0.01 | 24.43 ± 5.83 | 24.46 ± 5.83 | 0.07 ± 0.00 |
| K | *n.d.* | 1.29 ± 0.21 | 1.29 ± 0.21 | 0.01 ± 0.00 |
| D | *n.d.* | 0.37 ± 0.06 | 0.37 ± 0.06 | 0.00 ± 0.01 |
| E | *n.d.* | 0.55 ± 0.11 | 0.55 ± 0.11 | 0.01 ± 0.01 |
| S | *n.d.* | 0.90 ± 0.29 | 0.90 ± 0.29 | 0.02 ± 0.00 |
| T | *n.d.* | 1.04 ± 0.41 | 1.04 ± 0.41 | 0.02 ± 0.00 |
| N | *n.d.* | 0.78 ± 0.14 | 0.78 ± 0.14 | 0.00 ± 0.01 |
| Q | *n.d.* | 0.68 ± 0.10 | 0.68 ± 0.10 | 0.01 ± 0.01 |
| C | 0.07 ± 0.01 | 42.93 ± 12.84 | 43.00 ± 12.85 | 0.06 ± 0.01 |
| G | *n.d.* | 0.86 ± 0.18 | 0.86 ± 0.18 | 0.01 ± 0.00 |
| Y | *n.d.* | 18.21 ± 2.06 | 18.21 ± 2.06 | 0.05 ± 0.00 |
| P | *n.d.* | 7.20 ± 0.69 | 7.20 ± 0.69 | 0.02 ± 0.01 |
| A | *n.d.* | 3.44 ± 1.03 | 3.44 ± 1.03 | 0.02 ± 0.01 |
| V | *n.d.* | 4.30 ± 0.26 | 4.30 ± 0.26 | 0.03 ± 0.02 |
| I | *n.d.* | 2.70 ± 0.88 | 2.70 ± 0.88 | 0.04 ± 0.02 |
| L | *n.d.* | 2.32 ± 0.68 | 2.32 ± 0.68 | 0.02 ± 0.01 |
| M | *n.d.* | 8.44 ± 1.12 | 8.44 ± 1.12 | 0.02 ± 0.01 |
| F (WT) | 0.05 ± 0.03 | 32.55 ± 5.48 | 32.61 ± 5.51 | 0.09 ± 0.03 |
| W | 0.23 ± 0.04 | 159.26 ± 24.12 | 159.49 ± 24.16 | 0.29 ± 0.02 |

**Supplementary Table S5.** Butyl acetate titers measured by the colorimetric assay and GC/MS. $\triangle Ab_{520} = Ab_{520,AAT^+,BuOH^+} - Ab_{520,AAT^+,BuOH^-}$. $Ab_{520,AAT^+,BuOH^+} - Ab_{520,AAT^+,BuOH^-}$ represent the absorbances from cell cultures with or without an exogenous addition of 2 g/L butanol (BuOH), respectively. WT, wild type. Positive hits in bold.

| $ATF1_{Sc}$ variants | Colorimetric assay | | | | GC/MS |
|---|---|---|---|---|---|
| | $Ab_{520}^{BtOH}$ | $Ab_{520}^{None}$ | $\triangle Ab_{520}^{adj.}$ | Butyl acetate (mg/L) using $\triangle Ab_{520}^{adj.}$ | Butyl acetate (mg/L) |
| WT | 0.14 ± 0.01 | 0.11 ± 0.00 | 0.03 ± 0.01 | 20.33 ± 5.29 | 20.47 ± 1.84 |
| P348W | 0.14 ± 0.01 | 0.11 ± 0.00 | 0.03 ± 0.01 | 18.00 ± 8.98 | 21.29 ± 7.19 |
| P348R | 0.12 ± 0.00 | 0.11 ± 0.00 | 0.02 ± 0.00 | 11.33 ± 3.13 | 12.11 ± 5.38 |
| **P348M** | 0.22 ± 0.00 | 0.12 ± 0.01 | 0.10 ± 0.01 | **68.00 ± 6.93** | **69.88 ± 8.70** |
| P348H | 0.12 ± 0.00 | 0.12 ± 0.00 | 0.00 ± 0.01 | 0.00 ± 3.38 | 5.34 ± 0.85 |
| P348K | 0.12 ± 0.00 | 0.11 ± 0.01 | 0.01 ± 0.01 | 8.00 ± 5.31 | 4.43 ± 1.01 |
| P348N | 0.12 ± 0.00 | 0.12 ± 0.00 | 0.00 ± 0.00 | 0.17 ± 1.99 | 5.01 ± 1.67 |
| P348I | 0.13 ± 0.00 | 0.11 ± 0.00 | 0.02 ± 0.00 | 13.00 ± 1.68 | 7.93 ± 2.94 |
| **P348S** | 0.20 ± 0.01 | 0.11 ± 0.00 | 0.09 ± 0.01 | **58.50 ± 7.59** | **62.66 ± 19.60** |
| P348D | 0.12 ± 0.00 | 0.11 ± 0.00 | 0.01 ± 0.00 | 9.17 ± 2.40 | 6.88 ± 1.51 |
| P348C | 0.14 ± 0.00 | 0.11 ± 0.00 | 0.03 ± 0.00 | 19.17 ± 1.14 | 20.48 ± 4.43 |
| **P348A** | 0.20 ± 0.02 | 0.11 ± 0.00 | 0.09 ± 0.02 | **59.00 ± 12.90** | **52.54 ± 22.65** |
| P348Q | 0.13 ± 0.00 | 0.10 ± 0.00 | 0.03 ± 0.00 | 17.67 ± 2.80 | 20.25 ± 4.98 |

**Supplementary Figure S1.** Growth kinetics of the BL21 (DE3) strains carrying the 20 $CAT_{Sa}$ F97 variants in microplates w/ or w/o hexadecane overlay. Blue square indicates w/o overlay; Orange circle indicates w/ overlay.

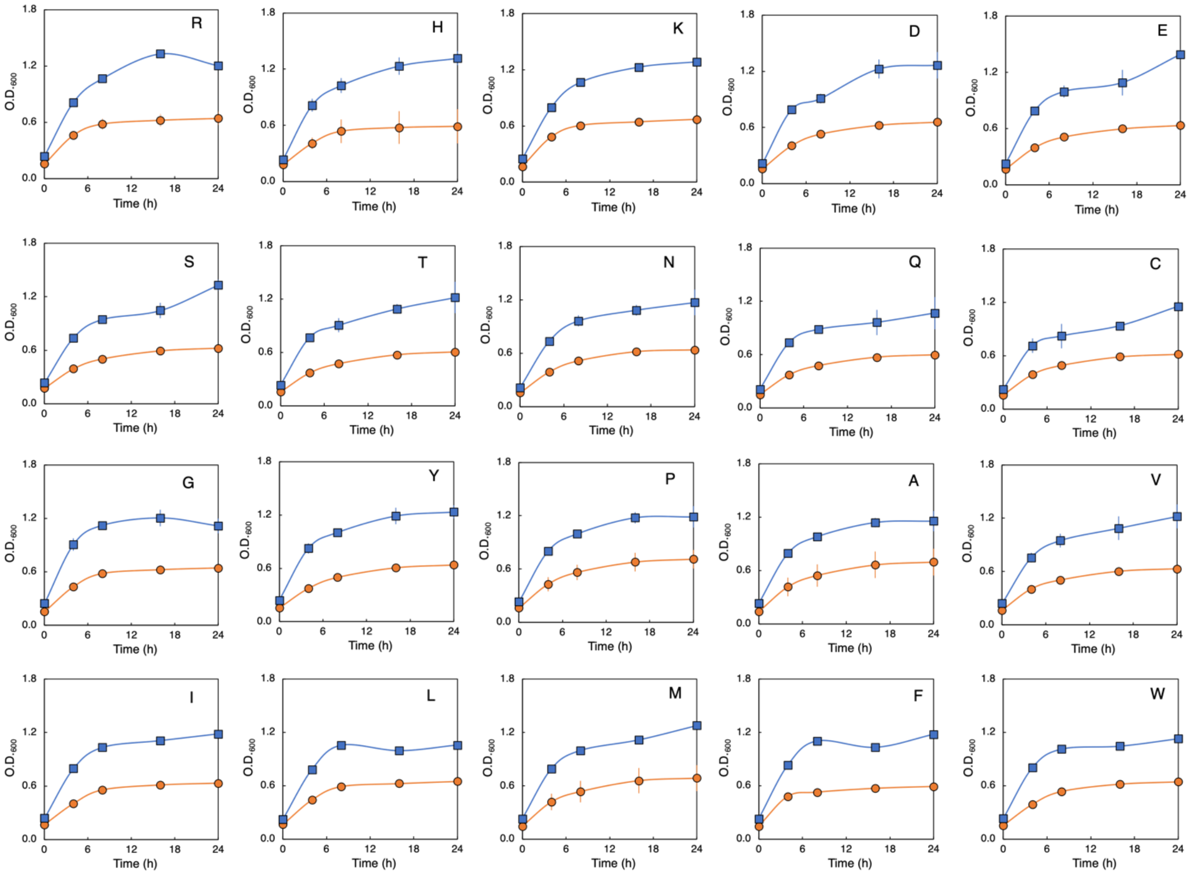

**Supplementary Figure S2.** (**A**) Two competing pathways for microbial biosynthesis of acetate esters with isobutanol doping. Filled line represents the ester synthesis pathway by $CAT_{Sa}$. Dotted lines represent the endogenous ethanol pathway causing the synthesis of inevitable EA byproduct. (**B**) Comparison of the IBA titers measured by the high-throughput microbial screening and GC/MS methods. For the colorimetric assay, baseline subtraction from cultures without alcohol doping is important to accurately estimate ester production. The error bars represent standard deviation of three biological replicates (*n*=3).

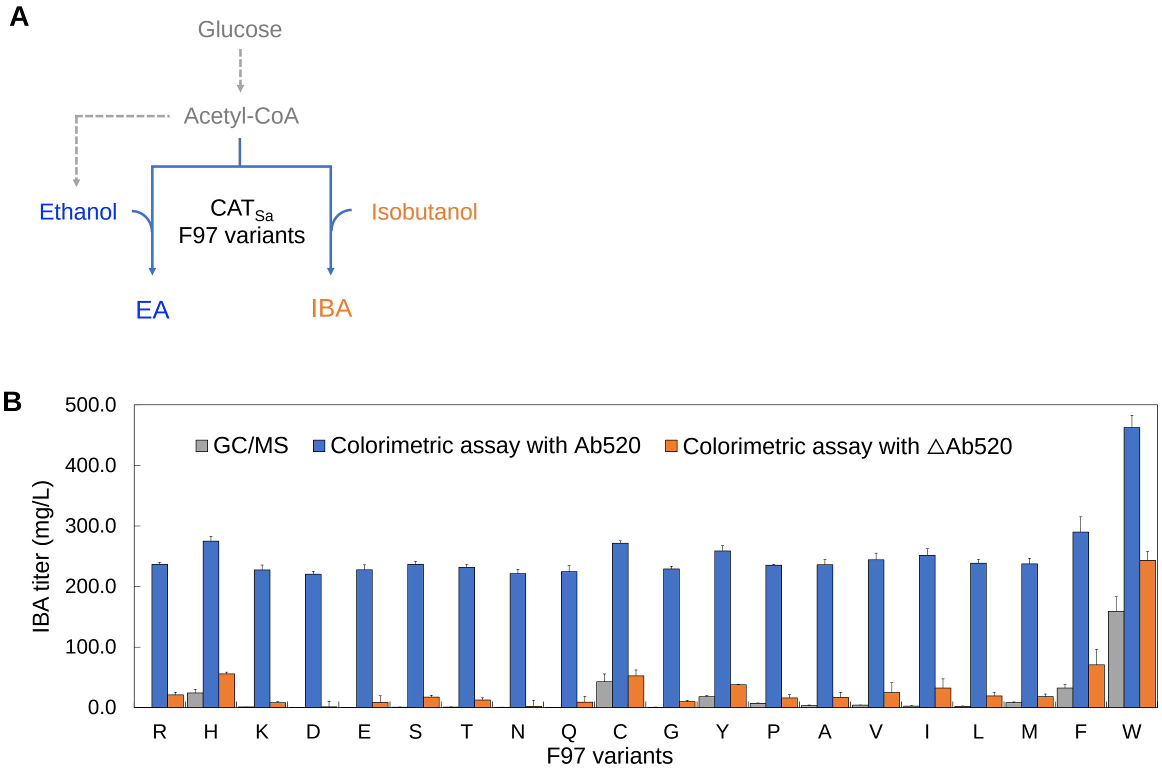

**Supplementary Figure S3. (A)** The 3D structure of homology model of ATF1$_{Sc}$. The binding pocket of ATF1$_{Sc}$ is shown in a yellow cloud while binding pocket residues of ATF1$_{Sc}$ appear in blue. **(B)** The 3D structure of acetyl-CoA-butanol-ATF1$_{Sc}$ complex. The residues interacting with butanol in the acetyl-CoA-butanol-ATF1$_{Sc}$ complex are in green.

**A**

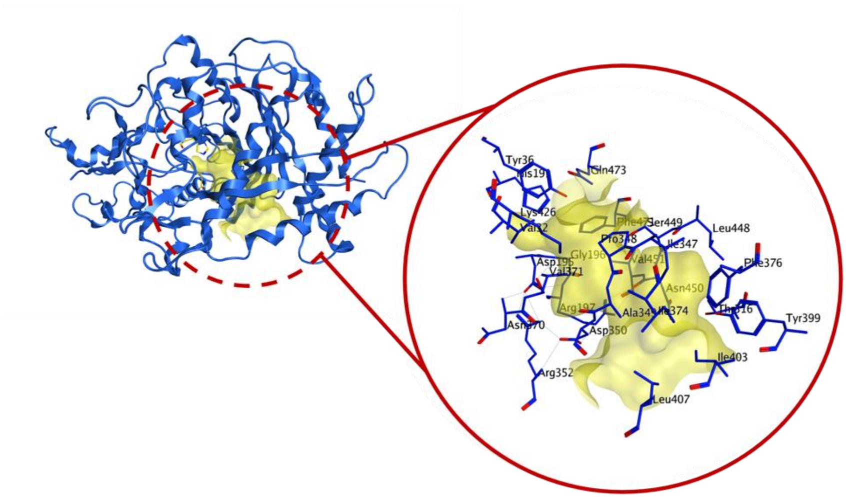


Identified the binding pocket residues

| Entry | Residues |
|---|---|
| 1 | V32 |
| 2 | Y36 |
| 3 | H191 |
| 4 | D195 |
| 5 | G196 |
| 6 | R197 |
| 7 | T316 |
| 8 | I347 |
| 9 | P348 |
| 10 | A349 |
| 11 | D350 |
| 12 | R352 |
| 13 | N370 |
| 14 | V371 |
| 15 | I374 |
| 16 | F376 |
| 17 | Y399 |
| 18 | I403 |
| 19 | L407 |
| 20 | K426 |
| 21 | L448 |
| 22 | S449 |
| 23 | N450 |
| 24 | V451 |
| 25 | F471 |
| 26 | Q473 |

**B**

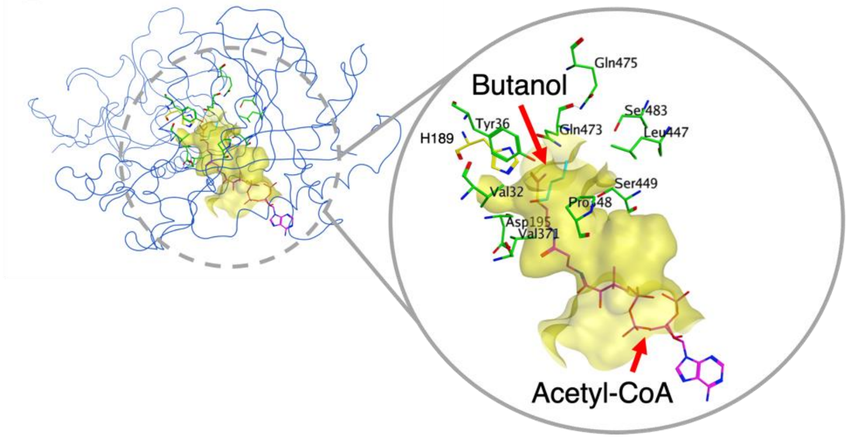


Residues interacting with butanol

| Entry | Residues |
|---|---|
| 1 | V32 |
| 2 | Y36 |
| 3 | D195 |
| 4 | P348 |
| 5 | V371 |
| 6 | L447 |
| 7 | S449 |
| 8 | Q473 |
| 9 | Q475 |
| 10 | S483 |
